## Supplementary material for "Targeting Host PIM Protein Kinases Reduces Mayaro Virus Replication"

**Figure legends**

**Figure S1. PIM kinase inhibitors partially prevent MAYV-induced cytopathic effects.** HeLa cells or HDFs were pre-treated with PIM1 Inh 2, AZD1208 or DMSO (as a control) for 2 h. Then, the compounds were removed and the cells were infected with MAYV at an MOI of 1 (HDFs) or 10 (HeLa cells). After 1 h of virus absorption, virus was removed and PIM kinase inhibitors or DMSO were added to the cells in fresh medium and incubated for an additional 48 h. Next, cytopathic effects were evaluated with an inverted microscope and an MCI70-HD camera (Leica). Scale bar: 100 µm.

**Figure S2. PIM kinase inhibitors have no virucidal effect on MAYV.** 1 x $10^7$ PFU of MAYV were incubated with PIM kinase inhibitors in serum-free medium for 2 h at 37 °C. Next, the virus was directly titrated using a plaque-forming assay. Data were analyzed with the Mann & Whitney test. Statistically significant differences are denoted as follows: ns: non-significant.

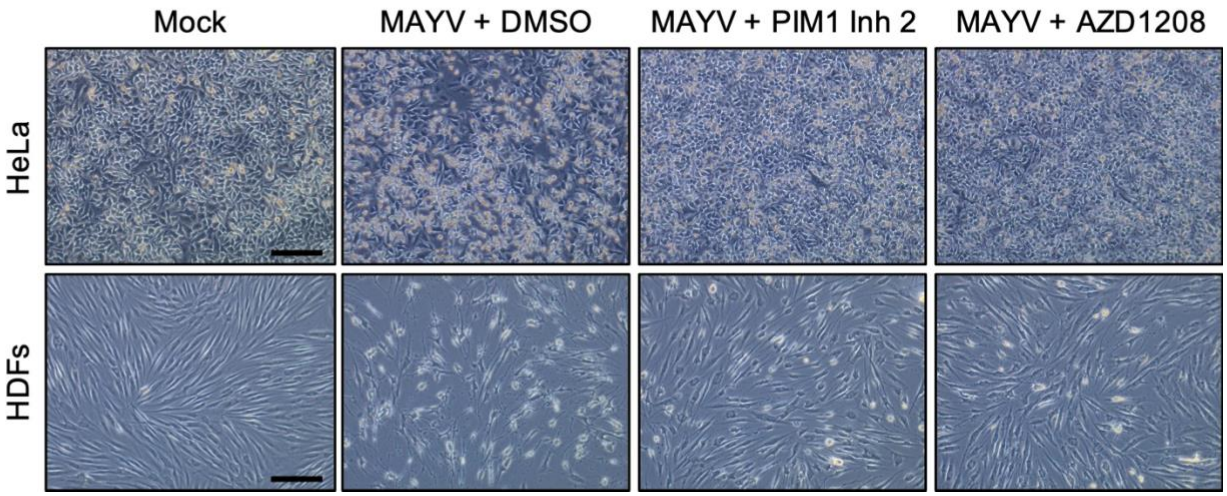


**Figure S1**

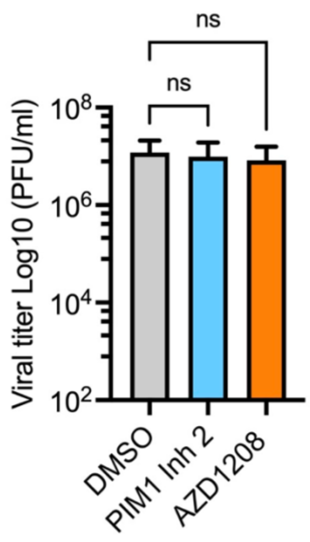
ns
ns
Viral titer Log10 (PFU/ml)
$10^8$
$10^6$
$10^4$
$10^2$
DMSO
PIM1 Inh 2
AZD1208

**Figure S2**
